# Human long noncoding RNA, *VILMIR,* is induced by major respiratory viral infections and modulates the host interferon response

**DOI:** 10.1101/2024.03.25.586578

**Authors:** Kristen John, Ian Huntress, Ethan Smith, Hsuan Chou, Tammy S. Tollison, Sergio Covarrubias, Elisa Crisci, Susan Carpenter, Xinxia Peng

## Abstract

Long noncoding RNAs (lncRNAs) are a newer class of noncoding transcripts identified as key regulators of biological processes. Here we aimed to identify novel lncRNA targets that play critical roles in major human respiratory viral infections by systematically mining large-scale transcriptomic datasets. Using bulk RNA-sequencing (RNA-seq) analysis, we identified a previously uncharacterized lncRNA, named virus inducible lncRNA modulator of interferon response (*VILMIR)*, that was consistently upregulated after *in vitro* influenza infection across multiple human epithelial cell lines and influenza A virus subtypes. *VILMIR* was also upregulated after SARS-CoV-2 and RSV infections *in vitro*. We experimentally confirmed the response of *VILMIR* to influenza infection and interferon-beta (IFN-β) treatment in the A549 human epithelial cell line and found the expression of *VILMIR* was robustly induced by IFN-β treatment in a dose and time-specific manner. Single cell RNA-seq analysis of bronchoalveolar lavage fluid (BALF) samples from COVID-19 patients uncovered that *VILMIR* was upregulated across various cell types including at least five immune cells. The upregulation of *VILMIR* in immune cells was further confirmed in the human T cell and monocyte cell lines, SUP-T1 and THP-1, after IFN-β treatment. Finally, we found that knockdown of *VILMIR* expression reduced the magnitude of host transcriptional responses to IFN-β treatment in A549 cells. Together, our results show that *VILMIR* is a novel interferon-stimulated gene (ISG) that regulates the host interferon response and may be a potential therapeutic target for human respiratory viral infections upon further mechanistic investigation.

**IMPORTANCE:** Identifying host factors that regulate the immune response to human respiratory viral infection is critical to developing new therapeutics. Human long noncoding RNAs (lncRNAs) have been found to play key regulatory roles during biological processes, however the majority of lncRNA functions within the host antiviral response remain unknown. In this study, we identified that a previously uncharacterized lncRNA, *VILMIR*, is upregulated after major respiratory viral infections including influenza, SARS-CoV-2, and RSV. We demonstrated that *VILMIR* is an interferon-stimulated gene that is upregulated after interferon-beta (IFN-β) in several human cell types. We also found that knockdown of *VILMIR* reduced the magnitude of host transcriptional responses to IFN-β treatment in human epithelial cells. Our results reveal that *VILMIR* regulates the host interferon response and may present a new therapeutic target during human respiratory viral infections.

## INTRODUCTION

With the first draft of the human genome researchers discovered that protein-coding genes only account for less than 2% of the genome (1). At first glance, the remaining 98% of intergenic sequence was considered “junk DNA.” However, advances in next generation sequencing technology revealed that about 80% of the genome may be transcribed, the majority of which is noncoding RNAs (ncRNAs) (2). While some ncRNAs are well characterized, such as microRNAs, much less is known about the function of long noncoding RNAs (lncRNAs). LncRNAs are transcripts greater than 500 nucleotides in length that have low translational potential (3). They are the largest class of ncRNAs with 20,424 human lncRNA genes annotated in the recent GENCODE V45 release (4), however identifying their functions has been challenging due to their low expression levels and low evolutionary conservation at the sequence level (5, 6). Despite these challenges, lncRNAs have emerged as key regulators of biological processes, such as transcription, mRNA stabilization, and protein translation (7). More recently, multiple lncRNAs have been identified in host antiviral and immune responses (8–10).

LncRNAs were first suggested to regulate viral infections when they were differentially expressed in mice after both severe acute respiratory syndrome coronavirus (SARS-CoV) and influenza infections (11). While RNA-sequencing (RNA-seq) analysis identified over 5,000 lncRNAs related to these viral infections, their specific roles in infection were not explored (12). Since then, individual lncRNAs have been studied in more detail and found to significantly alter influenza infection in human epithelial cells by either inhibiting or promoting viral infection (13). For example, lncRNA lnc-MxA was found to promote influenza A virus (IAV) infection by forming an RNA-DNA triplex at the interferon-beta (*IFN-β*) promoter and inhibiting the transcription of *IFN-β* (14). Lnc-ISG20 was found to inhibit IAV infection by competitively binding to microRNA 326 and reducing its inhibition of ISG20 translation (15). While a few IFN-independent lncRNAs have been identified in influenza infection (16–19), many of the identified lncRNAs within influenza infection are regulated by the IFN pathway to either promote the host immune response or control the immune response through negative feedback mechanisms (13).

In this study, we mined large-scale public RNA-seq data and identified a previously uncharacterized human lncRNA that we found was consistently upregulated after IAV infection across multiple human epithelial cell lines and influenza A virus subtypes. We showed that this lncRNA, *VILMIR*, was also upregulated after SARS-CoV-2 infection, RSV infection, and after IFN-β treatment. In addition, we analyzed single cell RNA-seq data from bronchoalveolar lavage fluid (BALF) samples from COVID-19 patients and demonstrated that *VILMIR* was upregulated in multiple cell types including epithelial cells and at least five immune cell types. Finally, we found that knockdown of *VILMIR* in human epithelial A549 cells broadly dampened the host response to IFN-β treatment. Our results show that lncRNA *VILMIR* is a novel IFN-stimulated gene (ISG) responding to major respiratory viral infections and may play a broad role in antiviral and innate immunity requiring further mechanistic investigation.

## MATERIALS AND METHODS

### RNA-sequencing data analysis

From public influenza infection experiments made available through the Gene Expression Omnibus (GEO) (20, 21), we collected 121 bulk RNA-seq samples. These samples included 4 human epithelial cell lines from the respiratory tract and 10 different influenza strains, and they ranged from 1 to 48 hours post influenza infection (summarized in Table 1). Infected and control samples were selected from experiments GSE97949 (22), GSE75699 (23, 24), GSE89008 (25), GSE68673 (26), GSE61517 (27), and GSE104168 (28). Samples were mapped using STAR version 2.5.2b (29) to the human genome assembly 38 (Hg38) with GENCODE V25. Custom STAR parameters were set as follows: twopassMode: Basic and alignIntronMax: 100000; otherwise, default STAR parameters were used. Raw gene counts were filtered, requiring at least one sample with greater than 50 reads. Remaining gene counts were normalized using the voom quantile normalization method (30) from the R package Limma (31). Each unique infected condition was compared to the time, strain, and cell type matched controls defined in their public experiment, for a total of 32 contrasts or comparisons. A gene was considered significantly differentially expressed for a comparison if its unadjusted p-value was less than 0.05 and the absolute value of the log2 fold change was greater than 0.58, i.e. a minimum change of 1.5-fold.

**Table 1.** Summary of the collection of RNA-seq data from influenza infections in human epithelial cells as seen in Figure 1.

| Epithelial cell type | IAV subtype | Strain | Abbr. | MOI | Time points | Ref. | Notes |
| --- | --- | --- | --- | --- | --- | --- | --- |
| Human tracheobronchial epithelial (HTBE) cells | H1N1 | A/California/04/09 | CA09 | 5 | 5 | (25) | #: Mutant H5N1 as described in ref (74).<br>*: Unclear from (26) which one of three viruses was used: A/Wisconsin/33 (WSN), A/New Caledonia/20/99, or a recombinant virus containing the M, HA, and NA segments of WSN in the background of A/Victoria/3/75. |
|  | H3N2 | A/Wyoming/03/03 | WY03 |  | 4 |  |  |
|  | H5N1 | A/Vietnam/1203/04<sup>#</sup> | H5N1 |  | 4 |  |  |
| Primary normal human bronchial epithelial (NHBE) cells | H1N1 | A/Puerto Rico/8/1934 | PR8 | 1 | 2 | (23, 24) |  |
| BEAS-2B (epithelial cells from normal human lung epithelium) | H3N2 | A/Brisbane/10/07 | B07 | 1 | 3 | (27) |  |
|  | H3N2 | A/Perth/16/09 | P09 |  | 3 |  |  |
|  | H3N2 | A/Udorn/307/72 | Udorn |  | 3 |  |  |
| A549 (adenocarcinomic human alveolar basal epithelial cells) | H1N1<sup>*</sup> | A/Wisconsin/33 | <sup>*</sup> | 3 | 1 | (26) |  |
|  | H7N9 | A/Anhui/1/2013 | H7N9 | 1 | 3 | (22) |  |
|  | H1N1 | A/Puerto Rico/8/1934 | PR8 | 0.5 | 2 | (28) |  |
|  | H3N2 | A/New York/238/2005 | NY05 |  | 2 |  |  |

To select lncRNAs that were more relevant to IAV infection, we applied three major selection criteria. 1) Genes were consistently detected across all but one of the human epithelial cell lines (i.e. one count per million reads mapped (CPM) in all the uninfected control groups or one of the infection groups for at least one timepoint). 2) The same gene consistently exhibited large expression changes when the overall host response peaked during infection (upregulation or downregulation of 4-fold or more in at least 5 of 10 peak time points). 3) The same gene exhibited significant expression changes early after infection (6-7 hours after infection) with a fold change of 1.5 or more in at least 6 of 9 early time points (to account for smaller and more variable expression changes at earlier time points).

To investigate if *VILMIR* was differentially expressed under additional conditions, we separately analyzed 78 single-end RNA-seq samples (120 to 140 bp read length) from GSE147507 (32, 33). For this analysis, reads were mapped using STAR version 2.7.10b (29) to the human genome assembly 38 (Hg38) with GENCODE version 44. Lowly expressed genes were filtered out using the filterByExpr function in edgeR version 3.40.2 (34) with default parameters. Counts were normalized using the TMM normalization method via the calcNormFactors function in edgeR. Differentially expressed genes were identified using the voom method (30) of the Limma R package version 3.54.2 (31). To calculate gene log fold changes in response to infection, we adopted the same sample comparisons previously defined by Blanco-Melo et al. (32).

### Cell culture

Human lung epithelial A549 (CCL-185), SUP-T1 (CRL-1942), and THP-1 (TIB-202) cells were purchased from American Type Culture Collection (ATCC, Manassas, VA, USA). Madin-Darby Canine Kidney (MDCK) cells (London Line, FR-58) was obtained through the International Reagent Resource, Influenza Division, WHO Collaborating Center for Surveillance, Epidemiology and Control of Influenza, Centers for Disease Control and Prevention, Atlanta, GA, USA. Human embryonic kidney (HEK) epithelial 293FT cells were ordered from Invitrogen. A549 cell lines were maintained in F-12K media with 10% fetal bovine serum (FBS). MDCK cells were maintained in DMEM media plus 100 U/mL penicillin and 100 µg/mL streptomycin (P/S), 0.2% bovine albumin fraction V, 25 mM HEPES buffer, and 5% FBS. SUP-T1 cells were maintained in RPMI 1640 media with 1% glutamax, P/S, and 10% FBS. THP-1 cells were maintained in the same media as SUP-T1 except for an additional supplement of 0.05 mM 2-mercaptoethanol (BME). HEK 293FT cells were maintained in DMEM media plus 1% glutamax and 10% FBS. All cell lines were kept at 37°C in a 5% CO_2_ incubator.

### Influenza virus stocks

Influenza A/California/04/2009 (H1N1) virus was ordered from BEI Resources (NR-13658) and propagated in MDCK cells as previously described by the World Health Organization (35). Briefly, MDCK cells at about 95% confluency were washed twice with 1X Dulbecco’s phosphate buffered saline (DPBS) and once with serum-free media and then infected with viral inoculum at a multiplicity of infection (MOI) of 0.0001 in serum-free media for one hour at 37°C, 5% CO_2_. After one hour, additional serum-free media containing 0.2% bovine albumin fraction V and 2 μg/mL TPCK-trypsin was added, and the cells were incubated at 37°C, 5% CO_2_ until cytopathic effect (CPE) reached at least 75%. The cell culture supernatant was harvested, centrifuged at 500 x g for 10 minutes, and stored at -80°C.

Virus titer was determined by Tissue Culture Infectious Dose (TCID_50_) assay using an ELISA as described previously (36). Briefly, MDCK cells were incubated with ½-log dilutions of virus in serum-free media for 18 hours at 37°C, 5% CO_2_. The next day, the cells were washed with DPBS and fixed in cold 80% acetone for 10 minutes. Viral nucleoprotein (NP) was detected by ELISA as described using a 1/1,000 dilution of the anti-NP antibody (Millipore cat. #MAB8257) and a 1/2,000 dilution of the horseradish peroxidase-labeled goat anti-mouse IgG (SeraCare cat. #52200460).

Freshly prepared substrate (10 mg of σ-phenylenediamine dihydrochloride (Sigma-Aldrich, cat. # P8287) per 20 mL of 0.05 M phosphate citrate buffer, pH 5.0, containing 0.03% sodium perborate) was added to each well, and the reaction was stopped with an equal volume of 0.5 M sulfuric acid. Absorbance was measured at 490 nm using a Tecan Spark microplate reader and the TCID_50_ was calculated using the Reed-Muench method (37).

### Virus infection

A549 cells were seeded overnight at 150,000 cells per well in 12-well plates in 1.5 mL media to reach ∼80% confluency the next day. The cell monolayer was washed twice with 1X DPBS and then incubated with virus at MOI 0.1 or mock inoculum in serum-free media for one hour at 37°C, 5% CO_2_. After one hour, the inoculum was removed, and the cells were washed again with 1X DPBS. Additional serum-free media containing 0.2% bovine albumin fraction V and 0.5 μg/mL TPCK-trypsin was added, and the cells were incubated at 37°C, 5% CO_2_. At the indicated time points post infection, RNA was extracted for analysis following the TRIzol RNA isolation method (Invitrogen).

### Interferon treatment

A549 cells were seeded overnight at 150,000 cells per well in 12-well plates in 1.5 mL media to reach ∼80% confluency the next day. The media was then removed, and the cell monolayer was washed with 1X DPBS prior to interferon treatment. Cells were treated with fresh A549 media with or without human IFN-β recombinant protein (R&D Systems™ 8499IF010) at the indicated concentrations. Cells were harvested at the indicated time points after treatment following the TRIzol RNA isolation method (Invitrogen). SUP-T1 and THP-1 cells were seeded at 1 million cells per well in 12-well plates in 1 mL and incubated in media with or without human IFN-β recombinant protein at 10 ng/mL for six hours and then harvested by TRIzol.

### RNA isolation and quantitative PCR

Total RNA was isolated from cells following the TRIzol isolation method (Invitrogen) and quantified using Nanodrop spectrophotometry. One μg of RNA was reverse-transcribed into cDNA using the QuantiTect Reverse Transcription Kit (Qiagen) containing both oligo-dT and random primers.

Quantitative PCR (qPCR) was performed on the cDNA using PowerUp SYBR Green Master Mix (Applied Biosystems). Relative expression of the indicated RNAs was determined using the ΔΔCt method with *GAPDH* or *18S* as an endogenous control. Statistical analysis of significance was performed in JMP Pro 16 software. The primer sequences used in this study are as follows: *GAPDH* F: GGTATCGTGGAAGGACTCATGAC; *GAPDH* R: ATGCCAGTGAGCTTCCCGTTCAG (16); *18S* F: GAACGTCTGCCCTATCAACTTTC*; 18S* R: GATGTGGTAGCCGTTTCTCAG; *VILMIR* F: GCTCCACCCTGAAAGTC; *VILMIR* R: CTACACAGTGCTGAGGAAA; *IFN-β* F: GCTCTCCTGTTGTGCTTCTCCAC; *IFN-β* R: CAATAGTCTCATTCCAGCCAGTGC (38); *STAT1* F: ATGCTTGCTTGGATCAGCTG; *STAT1* R: TAGGGTCATGTTCGTAGGTG; *GPB1* F: CGCTCTTAAACTTCAGGAACAG; *GBP1* R: CGTCGTCTCATTTTCGTCTGG; *MALAT1* F: TCCCCACAAGCAACTTCTCT; *MALAT1* R: CCTCGACACCATCGTTACCT; *DANCR* F: CGGAGGTGGATTCTGTTAGG; *DANCR* R: TCGGTGTAGCAAGTCTGGTG; *RHOT1* F: GCTCTGGAGGATGTCAAGAATG; *RHOT1* R: CGTGTCTCCCTCTCTGGATAA; *LRRC37B* F: GGACCTGGAGCTTAGCATAAC; *LRRC37B* R: GTCCAATCTCTGTAGTGGGTTC.

### 5’ and 3’ Rapid Amplification of cDNA ends (RACE)

5’ RACE of lncRNA *VILMIR* was performed using the SMARTer RACE 5’/3’ Kit (Takara, USA) according to the manufacturer’s instructions. To ensure a high level of expression of *VILMIR*, RNA from IFN-β treated A549 cells was used for first-strand cDNA synthesis. 5’ RACE PCR was then performed using the 5’-RACE-ready cDNA and a gene-specific-primer (GSP) with the sequence GCTCACCACCTGTAATCCCAGTAT. The 5’ RACE product was cloned into the pRACE vector provided by the RACE kit and sequenced with Sanger sequencing.

As obtaining a 3’ RACE product using the SMARTer RACE kit was not successful, a different protocol was used for this purpose. cDNA for 3’ RACE was generated using RNA from IFN-β treated A549 cells using the Template Switch RT Enzyme Mix (NEB, #M0466) and anchored primers. PCR was performed in two rounds using a nested primer design to enrich for *VILMIR* with forward GSPs (1^st^ round: ACGGTTTGGCTGATGGAAGATG, 2^nd^ round: CTCTGTGCTTCTAAACTCACTA) and a reverse primer to the 3’ anchor sequence introduced in RT. PCR was performed using Titanium Taq (Takara, #639208) with an extension time of 90s for 30 (1^st^ round) or 20 (2^nd^ round) cycles of amplification. PCR products were purified by PAGE using an overnight elution in 0.1X TAE on an orbital shaker. Purified round 2 PCR products were cloned using the NEB PCR Cloning Kit (NEB, #E1202) and sequenced with Sanger sequencing.

### Isolation of cytoplasmic and nuclear RNAs

Cytoplasmic, nuclear, and total RNA fractions were prepared from A549 cells according to the RNA Subcellular Isolation Kit (Active Motif, Cat. #25501). cDNA was prepared as described above and qPCR was performed to analyze *VILMIR* expression in both cellular fractions. Total RNA was used for normalization to calculate the percentage of total RNA by the equation % of Input = 100 x [2^ (Ct total RNA – Ct RNA fraction)]. *MALAT1* and *DANCR* were used as nuclear and cytoplasmic lncRNA controls, respectively.

### Plasmids and Cloning

The dCas9-KRAB construct was ordered from Addgene (#89567). The GFP construct for overexpression was constructed from a pSico lentiviral backbone with a bidirectional minimal EF1A-minimal CMV promoter expressing T2A flanked genes: zeocin-resistant (Zeo) and Green Fluorescent protein (GFP). The gRNA construct was published previously (39). Guide RNA (gRNA) sequences targeting the transcription start site of *STAT1* and *VILMIR* were designed using the web-based tool, CRISPick by Broad Institute (40, 41). Two highly ranked gRNAs were selected for each gene. The sequences of the gRNAs are as follows: STAT1g1, GGTCGCCTCTGCTCGGTCTG; STAT1g2, GGAGGGGCTCGGCTGCACCG; VILMIRg1, GTCATGCGGAGGACAAGGAA; VILMIRg2, CCTCCGCATGACGCCCGTGC. The sequence of the control gRNA targeting GFP was TGGTGGGCTAGCGGATCTGA. Each pair of gRNA spacer forward/reverse oligos were annealed and ligated into the gRNA construct via the AarI site.

### dCas9-KRAB cell line construction and validation

To produce lentivirus, HEK-293FT cells were co-transfected with each construct described above and lentiviral vectors, psPAX2 (Addgene #12260) and pMD2.G (Addgene #12259) using the Lipofectamine 3000 Reagent (Invitrogen). A549 cells were first lentivirally infected with the dCas9-KRAB construct (described above) and were selected using blastocidin (10 µg/ml) for 1 week to obtain dCas9-KRAB-expressing cells. The same cells were lentivirally infected with the GFP construct and selected using zeocin (200-300 µg/ml) for 1 week. Next, the dCas9-KRAB-GFP-expressing cells were clonally expanded and infected with a GFP-targeting gRNA and selected with puromycin (2-5 µg/mL) for 1 week to establish efficient GFP knockdown using flow cytometry as a read out. *STAT1* and *VILMIR* gRNAs were then inserted into the most efficient dCas9-KRAB-GFP cells and selected to assay the effects of knockdown.

### Single cell RNA-seq data analysis

We collected nine SARS-CoV-2 infected 5’ single cell RNA-seq samples and three uninfected 5’ single cell RNA-seq samples from Liao et al. (GSE145926) (42). The Cell Ranger Software Suite version 7.0.1 (43) was used to align sequencing reads and generate feature-barcode matrices. Reads were mapped to the human genome assembly 38 (Hg38) with GENCODE V45 annotation provided by Ensembl (44). The Seurat R package version 4.3.0 (45) was used to join the cell type annotations from the final barcode-cell type mapping matrix published by Liao et al. (42) to the corresponding cells in the feature-barcode matrices produced by Cell Ranger. Cells that were not assigned an identity by Liao et al. (42) were excluded from downstream analysis. Additionally, cell types which had low relative abundance (<0.15% of annotated infected cells) in the infected samples were removed from the analysis. Cell types which were considered abundant in the infected samples but had low relative abundance in control samples (<0.15% of annotated control cells) were retained but had their control data excluded from the analysis. For cell types with sufficient cell counts in each infected and control samples (>50 cells), a Wilcoxon rank sum test was conducted comparing the percentages of *VILMIR* positive cells in infected vs uninfected samples, separately for each cell type.

### cDNA library construction, RNA-sequencing, and IPA analysis

mRNA-sequencing was performed in biological triplicates in A549 *STAT1* KD, *VILMIR* KD, and control cell lines treated with mock or either 1 ng/mL or 10 ng/mL human IFN-β for 6 hours. Total RNA was isolated from cells following the TRIzol isolation method (Invitrogen). All samples were quantified and assayed to confirm minimum RNA integrity number of at least 9.4 using an Agilent Bioanalyzer (RNA 6000 Pico Kit, catalog no. 5067-1513). Next, 500 ng of total RNA per sample underwent mRNA capture and was then fragmented at 94 °C for 6 min. Sequencing libraries were prepared according to the manufacturer’s protocol using eleven cycles of final amplification (KAPA mRNA HyperPrep Kit, catalog no. KK8580 and KAPA UDI Adapter Kit, catalog no. KK8727).

Libraries underwent QC prior to sequencing using an Agilent Bioanalyzer (High Sensitivity DNA Kit, catalog no. 5067-4626). Next-generation sequencing was performed on an Illumina NextSeq500 (75-bp paired end) to a targeted depth of ∼20 million reads per sample.

Illumina RNA-seq reads were mapped against the human genome assembly version 38 (Hg38) using STAR version 2.7.9a (29). Custom STAR parameters were set as follows: limitOutSAMoneReadBytes: 1000000, outSAMprimaryFlag: AllBestScore, outFilterType: BySJout, alignSJoverhangMin: 8, alignSJDBoverhangMin: 3, outFilterMismatchNmax: 999, alignIntronMin: 20, alignIntronMax: 1000000, alignMatesGapMax: 1000000, outFilterMultimapNmax: 20; otherwise, default STAR parameters were used. Following read mapping, a count matrix was generated from the STAR results using R. Genes were removed from the matrix if they did not have at least 30 reads in a minimum of 3 samples from either the knockdown or control group. Counts were normalized using the TMM normalization method via the calcNormFactors function in edgeR version 3.40.2 (34).

To conduct our differential gene expression analysis, we utilized the limma-trend approach from Limma version 3.54.2 (31). Our analysis probed the differential gene expression of each of the two interferon treatments for each gene knockdown (including the GFP control) against their respective mock treatments, resulting in a total of 8 contrasts. We then contrasted the differential expression results of each knockdown (STAT1g1/1ng IFN-β vs STAT1g1/Mock, VILMIRg1/1ng IFN-β vs VILMIRg1/Mock, etc.) against that of the corresponding control (Ctrl/1ng IFN-β vs Ctrl/Mock, Ctrl/10ng IFN-β vs Ctrl/Mock), resulting in a total of 6 additional contrasts. Significantly differentially expressed genes were defined using two different threshold criteria. For our first cutoff, genes were considered differentially expressed in a given contrast if their unadjusted p-value was less than 0.05 with no fold change requirement. Our second, stricter threshold, required an adjusted p-value of less than 0.05 with a minimum fold change of 1.25 (absolute value of log2 fold change > ∼0.32) for a gene to be considered differentially expressed. The results of our differential gene expression analysis were then plotted using the ComplexHeatmap R package version 2.14.0 (46). Loess curves of each contrast were plotted using the ggplot2 R package version 3.4.3 (47).

Pathway enrichment analysis and disease and function enrichment analysis were generated using QIAGEN Ingenuity Pathway Analysis (IPA) (48). A raw p-value cut-off of < 0.05 was used to define genes with significant expression changes after *VILMIR* KD in each cell line and IFN-β treatment. Canonical pathways analysis identified the pathways from the QIAGEN Ingenuity Pathway Analysis library of canonical pathways that were most significant to the data set, and the Diseases & Functions Analysis identified the biological functions and/or diseases that were most significant from the data set. Molecules from the data set that met the p-value cutoff of 0.05 (−log10 p-value 1.3) and were associated with a canonical pathway or function and/or disease in the QIAGEN Knowledge Base were considered for the analysis. A right-tailed Fisher’s Exact Test was used to calculate a p-value determining the probability that the association between the genes in the dataset and the canonical pathways or functions/disease is explained by chance alone.

## RESULTS

### Human lncRNA *VILMIR* is consistently upregulated after influenza infection by a compendium of bulk RNA-seq analysis

We started with identifying lncRNAs consistently differentially expressed during human influenza infections. To do so, we first compiled a large-scale RNA-seq compendium of influenza infections in four human epithelial cell lines including primary and immortalized cell lines and ten different IAV strains covering both seasonal IAV subtypes (H1N1 and H3N2) and highly pathogenic avian IAV viruses (H5N1 and H7N9), totaling 121 RNA-seq samples, summarized in Table 1 (22–28). To select lncRNAs that were more relevant to IAV infection, we searched for genes that were consistently differentially expressed across different infection conditions (described in Materials and Methods).

Briefly, we searched for genes that 1) were consistently detected across each of the human epithelial cell lines, 2) consistently exhibited large expression changes when the overall host response peaked during infection, and 3) exhibited significant expression changes early after infection. Applying these criteria to all genes, we obtained a list of 15 candidate ncRNA genes: 13 lncRNAs, one snoRNA, and one vault RNA, as well as 98 protein coding genes including those well known to be involved in influenza infection such as *ISG15*, *IRF7*, and *MX1*, validating our selection strategy. Here we report one of the candidate lncRNAs that met the criteria described above, ENSG00000277511 or *VILMIR* (virus inducible lncRNA modulator of interferon response), and showed robust upregulation after influenza infection across all RNA-seq conditions (Figure 1A), as well as after other respiratory viral infections and IFN-β treatment, detailed below. We reasoned that this robust transcriptional response after viral infection indicated that *VILMIR* may play an important functional role during viral infection.

**Figure 1.**
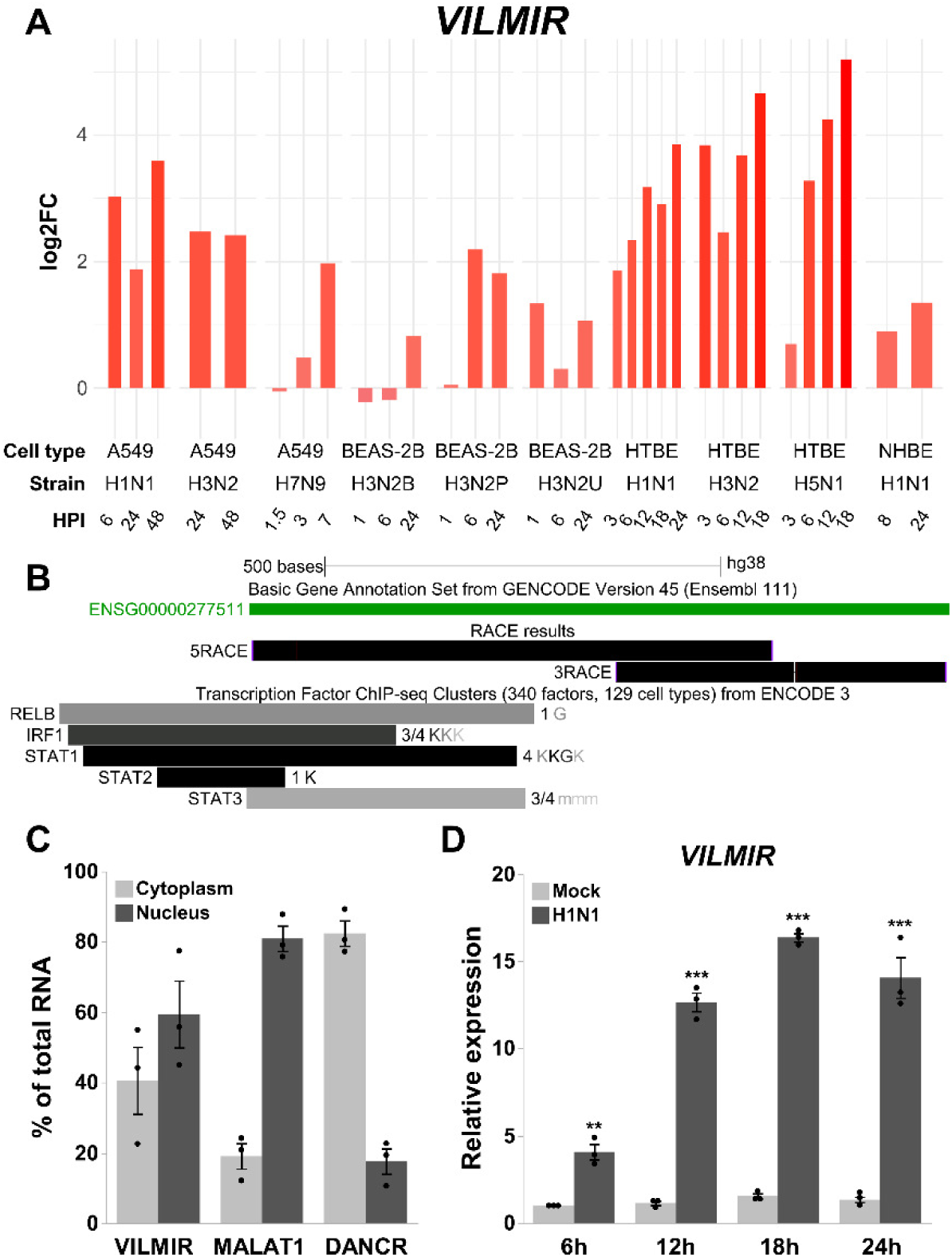
*VILMIR* identified as a lncRNA upregulated after influenza A infection in human epithelial cells. A) Temporal expression changes of lncRNA *VILMIR* in four human epithelial cell lines infected with various IAV strains in 121 RNA-seq samples (see Materials and Methods for data collection). The X-axis indicates human epithelial cell line, IAV strain, and hours post-infection (HPI). The Y-axis represents log2 fold change (log2FC) of *VILMIR* between infected samples and mock-treated control samples in each experiment. B) UCSC Genome Browser view of Sanger sequencing results of the 59 and 39 end sequences of lncRNA *VILMIR* determined by 59 and 39 RACE (black RACE results track) aligned to the Hg38 GENCODE V45 annotation (ENSG00000277511 in green). In addition, Transcription Factor ChIP-seq Clusters from ENCODE 3 showing RELB, IRF1, STAT1, STAT2, and STAT3 binding sites near the 59 end. Gray boxes represent the peak cluster of each transcription factor, and the darkness of the box is proportional to the maximum signal strength observed in any cell type. Labels to the right of the box represent cell types contributing to that cluster, with the darkness of the letters proportional to the signal strength in that cell line (ENCODE cell types are abbreviated as K = K562; G = GM12878; m = MCF_10A, MM.1S, mammary epithelial cell, medulloblastoma, myotube) (http://genome.ucsc.edu.). C) Expression of *VILMIR,* a nuclear control (*MALAT1*), and a cytoplasmic control (*DANCR*) was analyzed by RT-qPCR in cytoplasmic and nuclear fractions from A549 cells. Data was normalized to the total RNA fraction and expressed as % of total RNA (means ± SE; n=3). D) Relative expression of *VILMIR* was determined by RT-qPCR after IAV H1N1 CA/04/09 MOI 0.1 infection in A549 cells at the indicated time points and normalized to the mean of mock-treated cells at 6 hours. Data was normalized to 18S using the ΔΔCt method and expressed as means ± SE with individual replicates shown (n=3). *P < 0.05, **P < 0.01, ***P < 0.001 (Student9s t-test of H1N1 vs mock at each time point).

*VILMIR* is a long noncoding RNA transcript computationally annotated in Ensembl release 77 2014 (49). It is a single exon gene located on chromosome 17, 14 kilobases upstream of its closest neighboring protein-coding gene, *RHOT1,* and 74.083 kilobases downstream of its next closest protein-coding neighbor, *LRRC37B.* As the annotations of lncRNAs tend to be less accurate than that of protein-coding genes, we first validated the accuracy of the existing *VILMIR* annotation for its transcript boundaries, novel isoforms, and coding potential. We used 5’ and 3’ Rapid Amplification of cDNA Ends (RACE) in the human epithelial cell line A549 to confirm the Hg38 GENCODE V45 annotation of this gene (4). 5’ and 3’ RACE PCR fragments were Sanger sequenced and aligned to the GENCODE annotation in UCSC Genome Browser (50, 51). The analysis of 5’ and 3’ RACE products revealed that lncRNA *VILMIR* matches closely with the Hg38 GENCODE V45 annotation with a slightly shorter transcript of 877 nucleotides instead of 885, with no indication of additional novel isoforms (Figure 1B).

In addition to sequence, determining subcellular localization of lncRNA is also important in investigating function, as lncRNAs have been found to regulate transcription in the nucleus or translation and signaling in the cytoplasm, among other functions (52). We performed RNA fractionation in A549 cells and found that *VILMIR* co-localized in both nuclear and cytoplasmic fractions, suggesting it could function in or cycle between both compartments (Figure 1C). As *VILMIR* was co-localized in the cytoplasm, we investigated its coding potential by using the coding potential prediction tool, CPPRED (53). The tool used a support vector machine trained on known human coding and noncoding genes and reported only 1.07% probability that *VILMIR* had coding potential. In addition, since Ribosome-profiling (Ribo-seq) can experimentally uncover open reading frames (ORFs) within lncRNAs, we searched a recent publication cataloguing 7,264 ORFs found in lncRNAs and untranslated regions (UTRs) of protein-coding genes in Ribo-seq datasets from seven different publications (54), and found that *VILMIR* was not one of the identified lncRNAs that contains an ORF. These results indicate that *VILMIR* likely does not code for proteins.

Finally, as we relied on public RNA-seq data to prioritize *VILMIR* as a gene of interest, we sought to experimentally validate the response of *VILMIR* to influenza infection using RT-qPCR. We infected A549 epithelial cells with the commonly circulating H1N1 CA/09 IAV strain at a multiplicity of infection (MOI) of 0.1 and collected total RNA at various time points after infection to analyze expression. Our RT-qPCR results confirmed that *VILMIR* was significantly upregulated after H1N1 IAV infection at the earliest collected time point of 6 hours post-infection and increased up to 16-fold at 18 hours post-infection (Figure 1D). Expression of *VILMIR* began to plateau and drop at 24 hours post-infection, which is also when cytopathic effects on the cells were observed, indicating this drop is likely due to cell death.

### *VILMIR* expression is upregulated across multiple respiratory viral infections

To investigate if the upregulation of *VILMIR* was unique to influenza infection, we collected additional RNA-seq data (32, 33) and calculated *VILMIR* expression changes during several respiratory viral infections as well as IFN-β treatment in various human epithelial cell types including primary and immortalized cell lines. We observed that *VILMIR* was upregulated in infected or treated samples compared to mock-treated samples after RSV infection, SARS-CoV-2 infection, additional influenza infections, and IFN-β treatment (Figure 2A). As expected, the expression levels of genes in canonical interferon signaling such as interferon and JAK*/*STAT genes were also upregulated in these conditions (Figure 2B).

**Figure 2:**
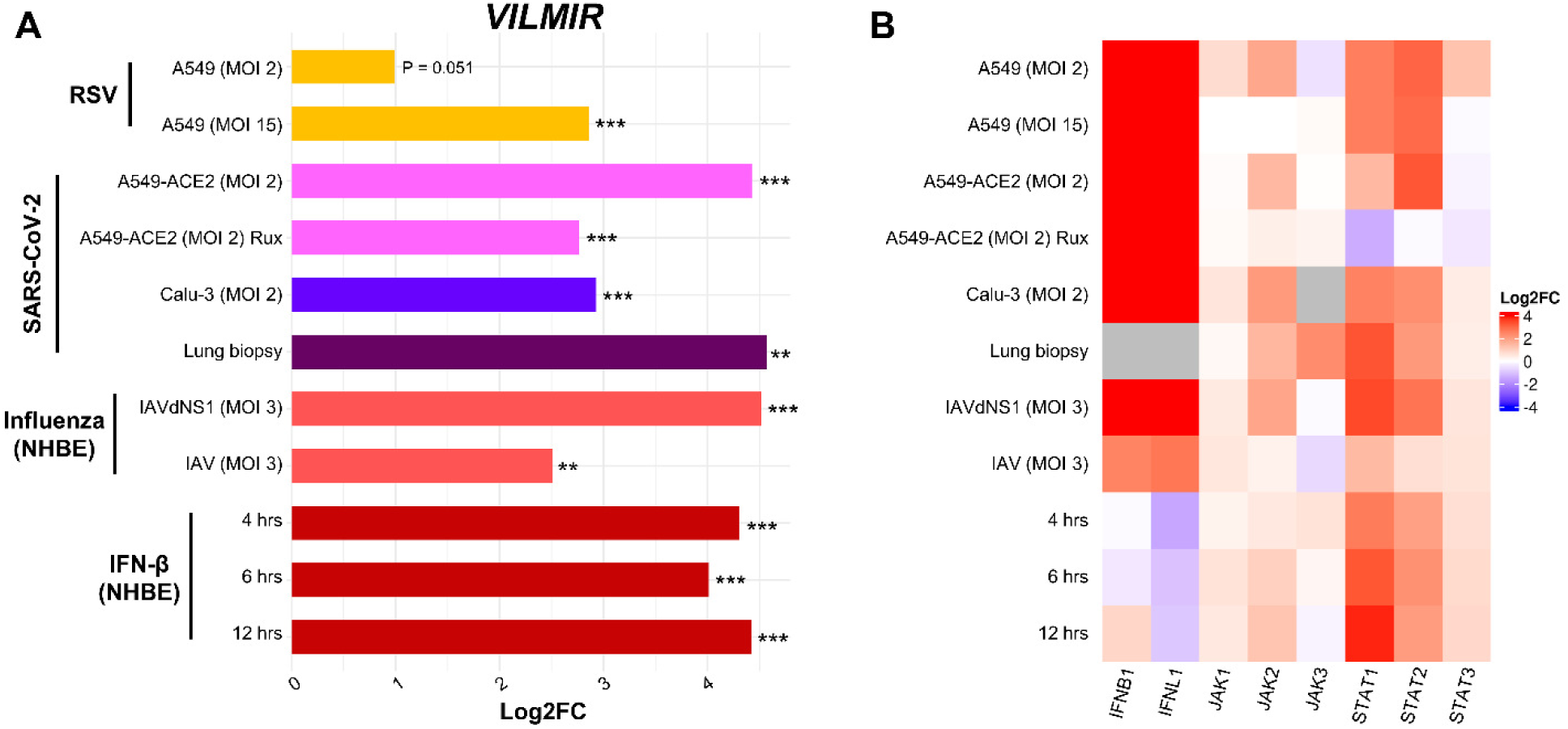
*VILMIR* is upregulated in response to RSV, SARS-CoV-2, influenza A viruses, and interferon-β. A) Horizontal barplot showing log2 fold change (Log2FC) between infected or treated samples and their corresponding mock-treated control samples in each condition as indicated by the label on the left. The Y-axis represents experimental conditions of RNA-seq samples from GSE147507 (32, 33). The color of the bars represents the virus and host cell-line for each set of samples. From top to bottom: yellow color for samples treated with Respiratory Syncytial Virus A2 strain; pink, blue, and purple for samples infected with Severe Acute Respiratory Syndrome Coronavirus 2 in the labeled cell types; salmon for samples treated with Influenza A/Puerto Rico/8/1934 WT or an NS1-deleted strain (IAVdNS1); and red for samples treated with human interferon-beta (IFN-β). B) Heatmap showing the expression changes (Log2FC) of selected *IFN*, *JAK* and *STAT* genes in the same samples when compared to matched controls. Red: upregulation; blue: downregulation; gray: insufficient numbers of RNA-seq reads to produce a measure. **P < 0.01, ***P < 0.001 (unadjusted p-value of infected or treated vs mock).

Interestingly, *VILMIR* upregulation decreased when host interferon responses were suppressed. For example, the upregulation of *VILMIR* decreased significantly (log2 fold change (log2FC) of 2 or a decrease of 4-fold, unadjusted p-value < 0.001) in Normal Human Bronchial Epithelial (NHBE) cells infected with wild-type IAV (log2FC 2.5, unadj. p-value < 0.01) compared to cells infected with a mutant IAV lacking the NS1 protein (dNS1) (log2FC 4.5, unadj. p-value < 0.001), a known inhibitor of the host interferon response (55) (Figure 2A). This also aligned with decreased upregulation of multiple IFN signaling genes in the wild-type IAV-infected cells, such as *IFNB1*, *IFNL1*, *STAT1*, and *STAT2* (Figure 2B). In SARS-CoV-2 infection of ACE2-modified A549 cells, the upregulation of *VILMIR* also decreased significantly (log2FC of 1.7 or a decrease of 3.2-fold, unadj. p-value < 0.001) in infected cells pretreated with Ruxolitinib (log2FC 2.7, unadj. p-value < 0.001), a JAK1 and JAK2 inhibitor, compared to infected cells not pretreated with Ruxolitinib (log2FC 4.4, unadj. p-value < 0.001) (Figure 2A). As *VILMIR* upregulation decreased after each of these conditions but was not completely abolished, these results suggest that *VILMIR* may be regulated by multiple pathways.

### *VILMIR* expression in A549 cells is induced by interferon-β treatment in a dose and time specific manner and is not impacted by *STAT1* knockdown

The observations that *VILMIR* was induced after several respiratory viral infections as well as interferon treatment (Figure 2A) indicated that *VILMIR* was likely an interferon-stimulated gene (ISG). To evaluate whether the upregulation of *VILMIR* to interferon was dependent on the dose of interferon treatment, we treated A549 cells with increasing concentrations of human IFN-β from 0-10 ng/mL for 6 hours and measured *VILMIR* expression by RT-qPCR. We observed a dose response of *VILMIR* with relative expression increasing as the IFN-β concentration increased (Figure 3A).

**Figure 3.**
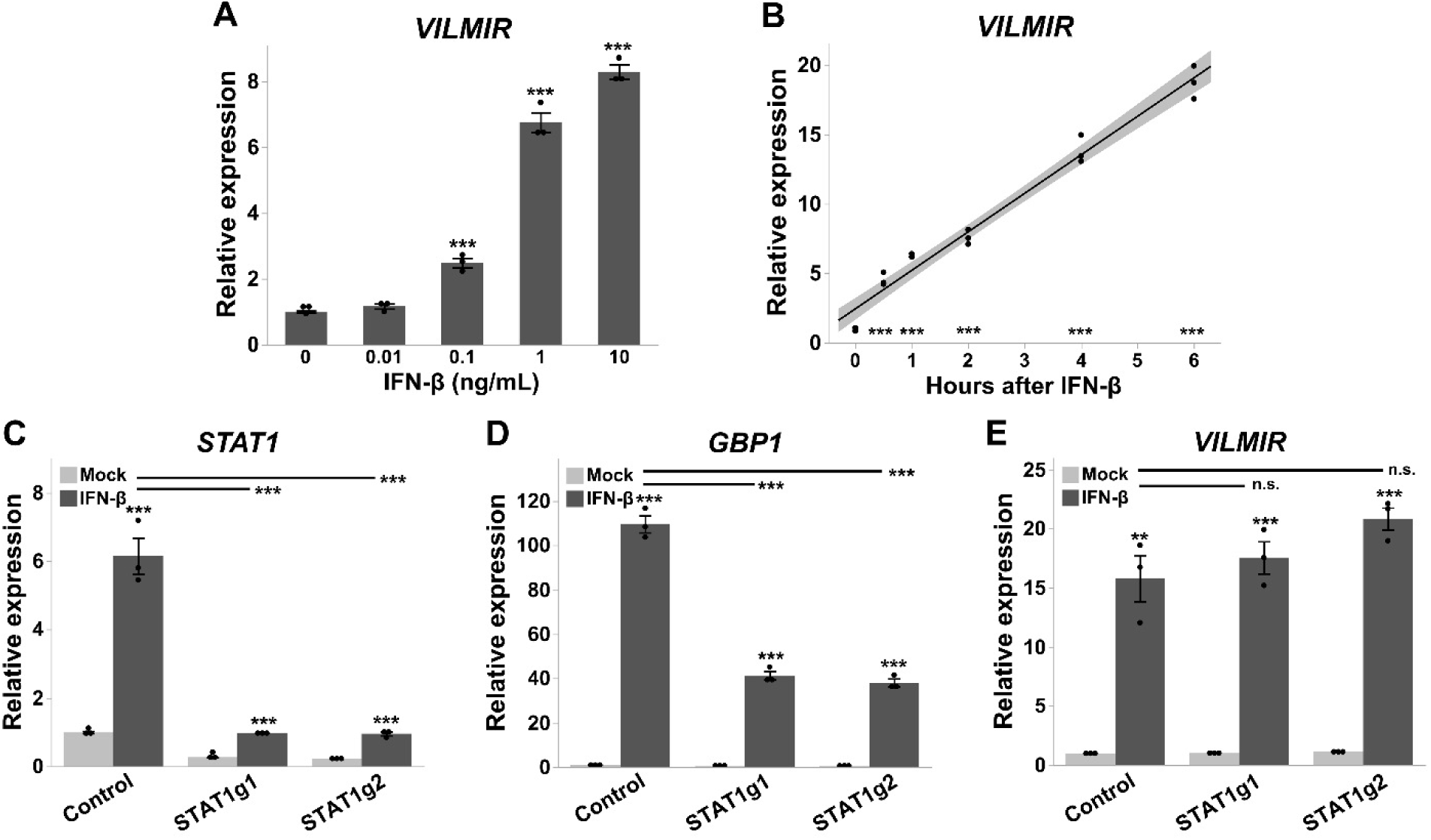
*VILMIR* expression after IFN-β treatment in A549 cells. A) Relative expression of *VILMIR* determined by RT-qPCR after treatment with the indicated concentrations of 0-10 ng/mL human IFN-β at 6 hours post-treatment in A549 cells (n=3). B) Relative expression of *VILMIR* determined by RT-qPCR after treatment with 10 ng/mL human IFN-β at 0, 0.5, 1, 2, 4, and 6 hours post-treatment in A549 cells. Individual replicates at each time point were plotted along with a regression line (black) and 95% confidence interval (gray shadow) (n=3, one-way ANOVA post hoc Dunnett). C-E) A549 cells expressing dCas9-KRAB were transduced with two guide RNAs targeting *STAT1* (STAT1g1 and STAT1g2) along with a control gRNA. The cell lines were treated with mock or 10 ng/mL human IFN-β for 6 hours. Relative expression of (C) *STAT1*, (D) *GBP1*, and (E) *VILMIR* in the gRNA lines was determined by RT-qPCR and normalized to the mean of the mock-treated control cell line. Unless otherwise stated, RT-qPCR data was normalized to GAPDH using the ΔΔCt method and expressed as means ± SE (n=3). *P < 0.05, **P < 0.01, ***P < 0.001 (Student9s t-test vs. 0 or the mock for each unless indicated).

We also found that *VILMIR* expression was significantly correlated with hours of IFN-β treatment, showing an increase in expression over a 6-hour time course and an immediate response to interferon within 30 minutes (Figure 3B). In addition, using Transcription Factor ChIP-seq Clusters from ENCODE 3 (56, 57), we found evidence for RELB (subunit of NF-κB), IRF1, STAT1, STAT2, and STAT3 binding sites near the 5’ end of *VILMIR* within several human cell lines (Figure 1B), further supporting our hypothesis that this gene may be a novel ISG.

To determine whether lncRNA *VILMIR* is directly induced by the canonical JAK/STAT signaling pathway or an independent pathway, the expression of *VILMIR* was analyzed by RT-qPCR after the knockdown of *STAT1* using a CRISPR interference system. Briefly, A549 cells expressing dCas9-KRAB and GFP were transduced with two different guide RNAs (gRNAs) targeting *STAT1* along with a negative control gRNA targeting GFP expression and treated with human IFN-β. As expected, in the negative control gRNA cell line, *STAT1* was upregulated after IFN-β treatment, whereas the STAT1g1 and STAT1g2 cell lines showed about an 84% knockdown of *STAT1* gene expression in the IFN-treated samples compared to the control cell line (Figure 3C). Successful knockdown of *STAT1* was also confirmed by the inhibition of known ISGs, such as *GBP1*, which had about a 65% decrease in expression compared to the control (Figure 3D).

Interestingly, there was not a significant difference in *VILMIR* expression despite knockdown of *STAT1* (Figure 3E). This indicates that *VILMIR* upregulation does not solely depend on *STAT1* transcription and may be regulated by an independent pathway or several pathways, as observed in the IAV dNS1 and SARS-CoV-2 infection analysis (Figure 2A).

### *VILMIR* is upregulated in multiple cell types during SARS-CoV-2 infection and interferon-β treatment

As our analysis of *VILMIR* was focused primarily on human epithelial cells, we were interested if *VILMIR* may be expressed in other cell types. We obtained published single cell RNA-seq data from bronchoalveolar lavage fluid (BALF) samples from COVID-19 patients and uninfected control samples (42). After confirming that *VILMIR* was detected in SARS-CoV-2 infected samples, we adopted the same cell type annotation as in (42) to investigate which cell types expressed *VILMIR*. We focused on a subset of nine annotated cell types that were relatively abundant in these samples (see Materials and Methods): macrophages, T cells, epithelial cells, natural killer cells (NK), neutrophils, monocyte derived dendritic cells (mDC), plasma cells, B cells, and plasmacytoid dendritic cells (pDC). Since for multiple cell types the number of cells in individual samples was zero or very low, for each cell type we calculated an overall percentage of cells where expression of *VILMIR* was detected by combining cells from individual samples, separately for infected and control samples (Figure 4A). Excluding three cell types (plasma, neutrophil, and pDC) which had extremely low numbers of cells in the uninfected control samples (15 cells in total for these three cell types, Table S1), we observed an increase in the overall percentage of cells expressing *VILMIR* in the SARS-CoV-2 infected samples when compared to uninfected control samples in the remaining six cell types (Figure 4A). We also performed the same calculation for *STAT1,* representing the canonical innate immune response and observed similar trends as expected (Figure 4B).

**Figure 4.**
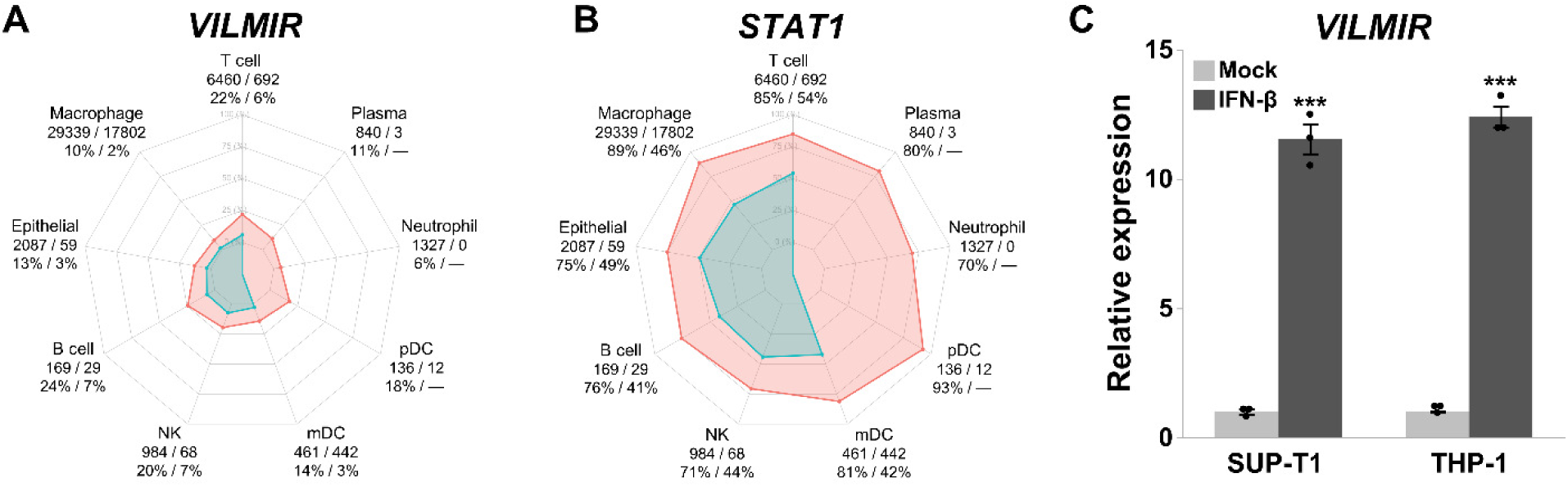
*VILMIR* is expressed in immune cell types after SARS-CoV-2 infection and IFN-β treatment. A) Overall percentage of cells expressing *VILMIR* from each cell type in SARS-CoV-2 infected and uninfected bronchoalveolar lavage fluid (BALF) from GSE145926 (42). Red points represent the overall percentage by combining cells of nine SARS-CoV-2 infected samples. Blue points represent the overall percentage by combining cells of three uninfected control samples. Each cell type is labeled with the total number of cells across infected and control samples, as well as the percentage of infected cells to control cells that express *VILMIR* (infected/control). A dash is indicated for cell types that had low relative abundance in control samples (see methods). Cell types are abbreviated as NK = natural killer; mDC = monocyte derived dendritic cells; and pDC = plasmacytoid dendritic cells. Sample-specific cell counts for each cell type can be found in Table S1. B) Similarly, as in A, the overall percentage of cells expressing *STAT1* from each cell type in SARS-CoV-2 infected and uninfected BALF. C) Relative expression of *VILMIR* determined by RT-qPCR after treatment with 10 ng/mL human IFN-β for 6 hours in the human cell lines, SUP-T1 T cells and THP-1 monocytes. All RT-qPCR data was normalized to GAPDH using the ΔΔCt method and expressed as means ± SE (n=3). ***P < 0.001 (Student9s t-test vs. the mock for each cell type).

For T cells and macrophages, the top two most abundant cell types that also had a sufficient number of cells in individual samples, we performed statistical testing and found the changes in percentage of cells expressing *VILMIR* to be statistically significant between the infected samples and the uninfected controls (Wilcoxon rank sum exact test unadjusted p-value < 0.01 for both).

To determine if upregulation of *VILMIR* could be related to the interferon response in T cells and macrophages, we treated the human T cell line SUP-T1 and the monocyte cell line THP-1 *in vitro* with 10 ng/mL human IFN-β for six hours. As expected, we observed significant upregulation of *VILMIR* in both cell lines after IFN-β treatment, confirming IFN-induced expression of *VILMIR* in human T cells and monocytes (Figure 4C). Overall, these results indicate that *VILMIR* can be activated in response to SARS-CoV-2 infection and IFN treatment in at least two immune cell types, in addition to epithelial cells as described above.

### Knockdown of *VILMIR* dampens the host response to interferon-β treatment in A549 epithelial cells

To investigate the functional role of *VILMIR* during the host interferon response, we repressed *VILMIR* expression using CRISPRi interference, as described above. A549 cells expressing dCas9-KRAB and GFP were transduced with two gRNAs targeting the 5’ end of *VILMIR* along with a negative control gRNA targeting GFP expression and treated with or without two separate concentrations of human IFN-β. Compared to the control cell line with a gRNA targeting GFP, we observed about 96% knockdown (KD) of *VILMIR* expression (Figure 5). We then performed RNA-seq analysis to identify the impact of *VILMIR* KD on the host transcriptional response to IFN-β treatment. In addition, a *STAT1* KD cell line (STAT1g1) was included for comparison, as *STAT1* is well-known to impact the host IFN response and served as a positive control for impact on host transcriptional response.

**Figure 5.**
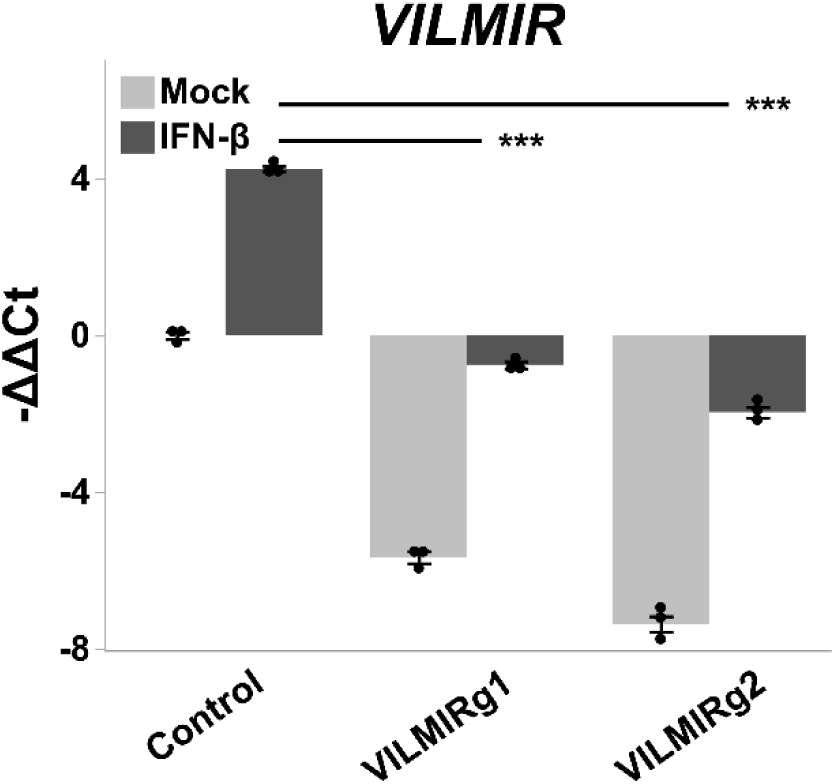
Knockdown efficiency of *VILMIR* expression after IFN-β treatment using CRISPRi. A549 cells expressing dCas9-KRAB were transduced with two gRNAs targeting *VILMIR* (VILMIRg1 and VILMIRg2) along with a control gRNA targeting GFP expression (Control) and treated with mock or 10 ng/mL human IFN-β for 6 hours. Relative expression of *VILMIR* was determined by RT-qPCR and normalized to the mean of the mock-treated control cell line. Data was normalized to GAPDH using the ΔΔCt method and expressed as means ± SE (n=3). ***P < 0.001 (Student9s t-test).

We first examined the overall effects of *VILMIR* KD on the expression responses to IFN-β treatment using a relatively relaxed criteria for differential expression analysis, i.e. raw p-value < 0.05, to investigate the potentially broad regulatory roles of *VILMIR*. Using this criterion, we identified 2,325 genes that showed altered expression changes to IFN-β treatment after *VILMIR* KD in at least one KD cell line treated with one of two doses of IFN-β (Figure 6A and Table S2). However, the average magnitude of expression changes was relatively small, with gene expression changing between 1.26-1.29-fold after *VILMIR* KD compared to the control. While *RHOT1* and *LRRC37B*, the two immediate neighboring protein-coding genes of *VILMIR*, were identified in this list of 2,325 genes, *RHOT1* expression changes were only significantly affected in one out of the four conditions.

**Figure 6.**
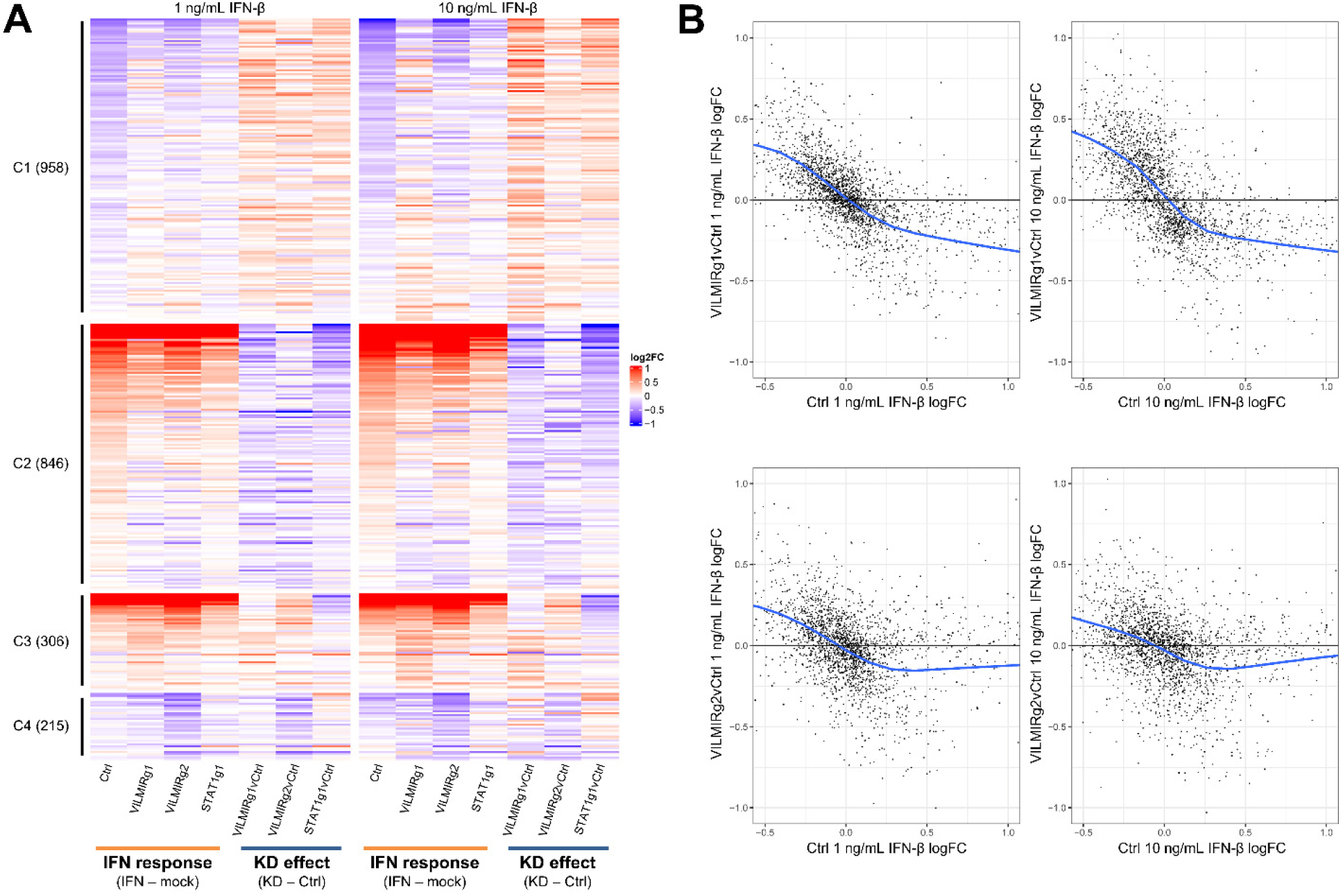
Transcriptional analysis of the effect of *VILMIR* knockdown on IFN-β treatment in A549 cells. A) Heatmap overview of the RNA-seq analysis of *VILMIR* knockdown (VILMIRg1 and VILMIRg2), *STAT1* knockdown (STAT1g1), or control (Ctrl) A549 gRNA cell lines that were treated with mock or either 1 ng/mL or 10 ng/mL human IFN-β for 6 hours (n=3). The heatmap displays 2,325 human genes that exhibited significant changes in their responses to IFN-β treatment after *VILMIR* knockdown (KD) in at least one KD cell line treated with one of two doses of IFN (raw p-value < 0.05). Rows are genes and columns are conditions and comparisons. As shown by the labels at the bottom, the log2 fold change (log2FC) after IFN-β treatment in each cell line was first calculated (<IFN response=), and then the <KD effect= was calculated by comparing the <IFN response= log2FC of each KD line to the <IFN response= log2FC of the control cell line. Red color indicates positive log2FC value (i.e. upregulation) in columns above the label 8IFN response9, or higher log2FC values in knockdown cells compared to that of control cells in columns above the label 8KD effect9. Blue color indicates negative log2FC value (i.e. downregulation) in columns above the label 8IFN response9, or lower log2FC values in knockdown cells compared to that of control cells in columns above the label 8KD effect9. Genes are grouped into four clusters based on their expression patterns across conditions/comparisons (columns): C1. The expression of 958 genes was downregulated by IFN treatment as shown by the 8Ctrl9 column but the magnitude of downregulation was decreased in *VILMIR* knockdown cells. C2. The expression of 846 genes was upregulated by IFN treatment as shown by the 8Ctrl9 column but the magnitude of upregulation was decreased in *VILMIR* knockdown cells. C3. The expression of 306 genes was upregulated by IFN treatment as shown by the 8Ctrl9 column but the magnitude of upregulation was increased in *VILMIR* knockdown cells. C4. The expression of 215 genes was downregulated by IFN treatment as shown by the 8Ctrl9 column but the magnitude of downregulation was increased in *VILMIR* knockdown cells. The full list of genes and log2FC values is available in Table S2. B) Scatterplots of the <KD effect=, i.e. the differences between log2FC values in each KD line to the log2FC values in control cell line, vs. the <IFN response= log2FC values in control cell line for the same 2,325 genes shown in (A). The blue line represents the locally estimated scatterplot smoothing (LOESS) curve and the black line at 0 on the y-axis represents if no change was observed after KD. Cut-offs were applied on the y-axis (−1.0-1.0) and x-axis (−0.5-1.0) for the data points. The full plots and mean squared error (MSE) can be seen in Figure S2.

Independent RT-qPCR analysis also showed the effect of *VILMIR* KD on the *RHOT1* expression changes was not significant (Figure S1A-C). In comparison, *LRRC37B* expression changes were significantly affected in three out of the four conditions by RNA-seq analysis, and this effect on *LRRC37B* expression changes was also confirmed by RT-qPCR for both *VILMIR* KD lines in the 10 ng/mL IFN-β treatment (Figure S1D-E). Since the observed magnitude of *VILMIR* KD effect on *LRRC37B* expression change was relatively small (average -ΔΔCt difference of -0.27 or a decrease of 0.83-fold by RT-qPCR) and *LRRC37B* was reported as a membrane receptor in neurons (58), the potential effect of *VILMIR* KD on *LRRC37B* needs to be further investigated.

To model the overall trend of the KD impact on host transcriptional response, a local regression analysis was performed by plotting the log2FC values in each KD line to the log2FC values in the control versus the log2FC values in the control cell line for the 2,325 genes identified in the RNA-seq data. Interestingly, we found that after *VILMIR* KD, the magnitudes of the expression changes induced by IFN-β treatment decreased in general, similarly as in the *STAT1* KD line (Figures 6B and S2). Generally, genes that were downregulated by IFN-β treatment also showed downregulation after *VILMIR* KD but with smaller fold changes (Figure 6A, Cluster 1). Likewise, genes that were upregulated by IFN-β treatment also showed upregulation after *VILMIR* KD but with smaller fold changes (Figure 6A, Cluster 2). In total, Clusters 1 and 2 represented 78% of the 2,325 genes identified. There were two smaller clusters of genes that were upregulated or downregulated with higher fold changes after *VILMIR* KD (Figure 6A, Clusters 3 and 4, respectively), representing 22% of the total genes identified. This indicates an overall opposite effect on host response between the KD of *VILMIR* and the IFN-β treatment. As KD of *STAT1* showed a similar trend in the same 2,325 genes, and *STAT1* is known to be a key activator of the IFN response, these results also suggest *VILMIR* might play an activating role in the IFN response.

To identify canonical pathways enriched in the differentially expressed genes impacted by *VILMIR* KD, QIAGEN Ingenuity Pathway Analysis (IPA) was performed (48). To account for differences in the dose of IFN-β, the two IFN-β treatments were separated for the pathway analysis. In the 1 ng/mL IFN-β treatment, 16 canonical pathways were significantly enriched in both KD lines with a raw enrichment p-value < 0.05 (-log10 p-value > 1.3, Figure 7A and Table S3), while 29 canonical pathways were significantly enriched in both KD lines in the 10 ng/mL IFN-β treatment (Figure 7B limited to top 17 pathways and Table S4). The longer list of 29 pathways enriched in the 10 ng/mL IFN-β treatment may be due to increased cellular stress caused by the higher dosage of IFN-β (59), as seven of the 29 pathways were specifically linked to cell death and survival functions, such as p53 Signaling, 14-3-3-mediated Signaling, and Death Receptor Signaling. Interestingly, the Interferon Signaling and JAK/STAT Signaling pathways were the only two pathways shared between both KD lines and IFN-β treatments and are both known to be important to the interferon response. Other relevant pathways to the interferon response and viral infection included TGF-β Signaling and the Senescence Pathway (cell cycle arrest) in the 1 ng/mL IFN-β treatment, as well as the Coronavirus Pathogenesis Pathway and ERK/MAPK Signaling in the 10 ng/mL IFN-β treatment.

**Figure 7.**
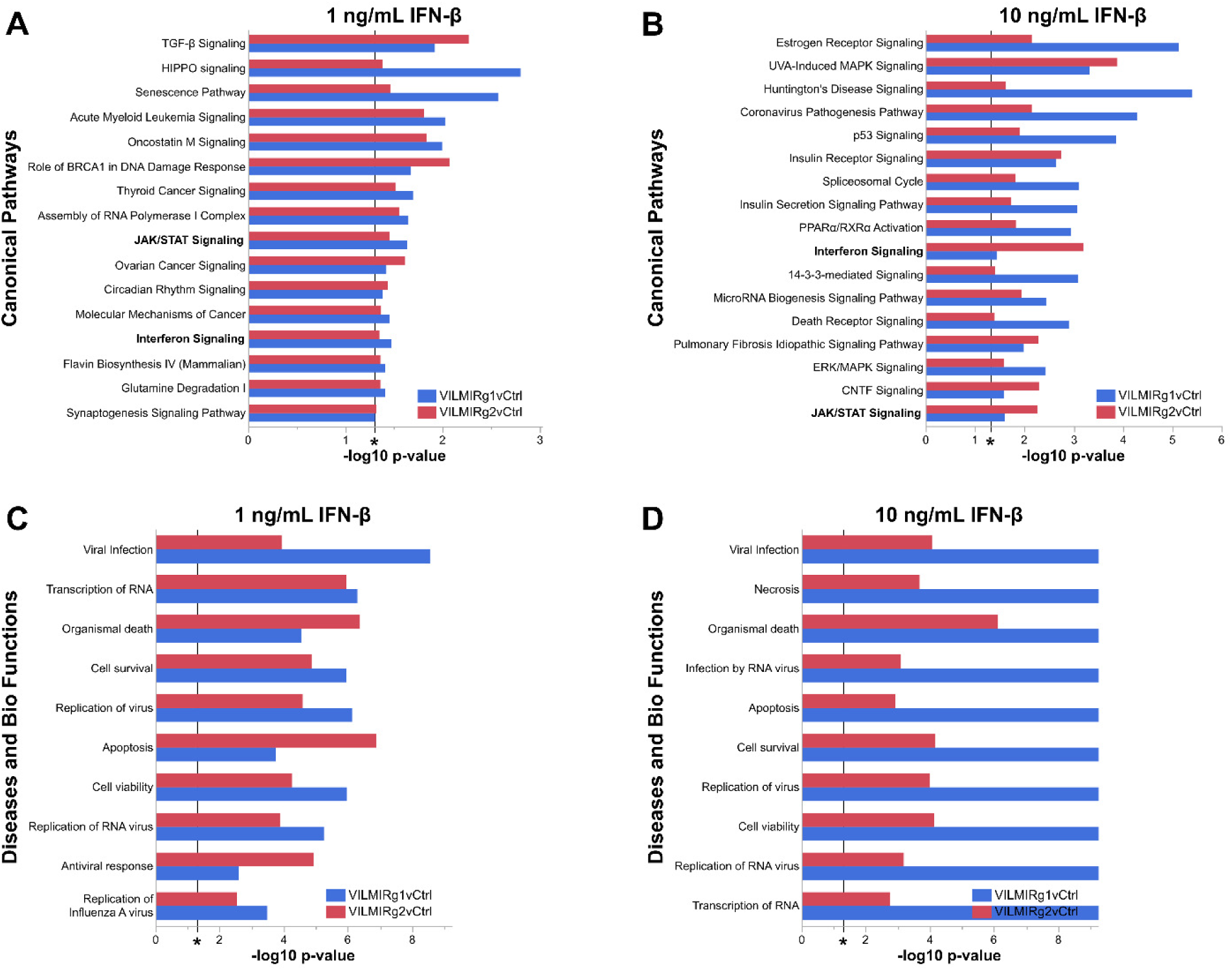
Pathways and functional categories enriched in differentially expressed genes induced by *VILMIR* knockdown after IFN-β treatment in A549 cells. A-B) QIAGEN Ingenuity Pathway Analysis (IPA) was performed to identify canonical pathways significantly enriched in the differentially expressed (DE) genes impacted by *VILMIR* KD after 1 ng/mL or 10 ng/mL IFN-β treatment in A549 cells. Enriched pathways shared between the IFN-β treatments are bolded. Panel B represents the top 17 out of 29 pathways. C-D) In addition, diseases and biological functions categories significantly enriched in the DE genes were identified for both IFN-β treatments using IPA (cancer categories excluded). Enriched pathways and functional categories that met the raw enrichment p-value < 0.05 (-log10 p-value cutoff of 1.3) using a right-tailed Fisher9s Exact Test in both *VILMIR* KD lines and were associated with a canonical pathway or function and/or disease in the QIAGEN Knowledge Base were included here (*P < 0.05 included as reference). For visualization, a -log10 p-value cutoff of 9.2 was applied to the x-axis of the bar plot in Panel D. Plots in B-D represent the subset of top enriched pathways and functional categories and the full lists are available in Tables S3-S6.

In addition, an enriched disease and biological function comparison was performed for differentially expressed genes impacted by *VILMIR* KD using IPA. While 141 diseases and functions classified by IPA were significantly represented in both KD lines and IFN-β treatments with a raw enrichment p-value < 0.05 (-log10 p-value > 1.3, Tables S5 and S6), this list decreased to 24 when cancer categories were excluded. Significantly enriched diseases and functions of potential relevance included viral infection, transcription of RNA, cell survival, replication of virus, and apoptosis (Figure 7C and D). It was observed that VILMIRg1 had much lower p-values than VILMIRg2 in the 10 ng/mL IFN-β treatment (Figure 7D), which is likely due to a difference in the number of DE genes in VILMIRg1 compared to VILMIRg2 (1,254 and 489 DE genes, respectively).

To examine the more robust expression changes after *VILMIR* knockdown and to mitigate potential off-target effects, we applied additional filtering to the differential expression analysis and obtained a list of 78 genes that showed altered expression changes to IFN-β treatment in the same direction across both KD cell lines by 1.25-fold or greater in at least one IFN-β treatment (Figure S3). The same observation was made for this subset of genes, with the majority of genes exhibiting smaller fold changes after *VILMIR* KD (Figure S3, Clusters 1 and 2). Again, a very small percentage of genes exhibited larger fold changes after *VILMIR* KD (Figure S3, Clusters 3 and 4). Two ISGs identified in this list of 78 were *IFIT2* and *IFI44L*, which decreased in their responses to IFN-β treatment after *VILMIR* KD. *IFIT2* decreased by 2-fold and 1.6-fold in the VILMIRg1 and VILMIRg2 KD lines, respectively, after 10 ng/mL IFN-β treatment. Similarly, *IFI44L* decreased by 1.5-fold and 1.8-fold in the VILMIRg1 and VILMIRg2 KD lines, respectively. *IFIT2* and *IFI44L* have previously been identified as ISGs during viral infection (60, 61), supporting our hypothesis that *VILMIR* may regulate the host interferon response. While the overall magnitude of observed changes after *VILMIR* KD was relatively small, these results indicate that *VILMIR* KD can broadly dampen the host response to interferon treatment in A549 epithelial cells.

## DISCUSSION

Here, through large scale RNA-seq analysis, we identified a previously uncharacterized human lncRNA, *VILMIR*, that is significantly upregulated in response to major respiratory viral infections such as influenza, SARS-CoV-2, and RSV in human epithelial cells. *VILMIR* is also upregulated in a dose and time-dependent manner after human IFN-β treatment. In addition, it responds to SARS-CoV-2 infection and IFN-β treatment in multiple immune cell types. Further, knockdown of *VILMIR* expression broadly dampens the host transcriptional response to IFN-β treatment in A549 cells. Together, these results indicate that *VILMIR* may broadly regulate the host interferon response, representing a potentially novel therapeutic target for modulating host response to infections.

Interestingly, as shown in RNA-seq analysis, *VILMIR* expression was reduced but not abolished in conditions with suppressed host IFN responses, such as after treatment with Ruxolitinib, a JAK1/2 inhibitor, or after infection with IAV with an intact NS1 protein. As expected, these conditions were also associated with decreased expression of IFN-signaling genes such as *IFNB1*, *IFNL1*, *STAT1*, and *STAT2*. The JAK proteins are phosphorylated upon binding of type I IFNs to IFN-α receptor 1 (IFNAR1) and IFNAR2, which then activate transcription factors such as STAT1 and STAT2 to induce transcription of ISGs that respond to viral infections (62). The influenza NS1 protein is a known antagonist of the host IFN response and has adapted multiple mechanisms to attenuate type I IFN production by disrupting cellular signaling (55). Other interferon-stimulated lncRNAs have been found to be affected by attenuation of the host immune response. For example, expression of the interferon-stimulated lncRNA *BISPR* was shown to be reduced after Ruxolitinib treatment in human epithelial cells as well as in cells with *STAT1* knockdown or inhibition of IRF1, suggesting that *BISPR* can respond to multiple transcription factors during the IFN response (63). Additionally, *BISPR* had increased expression in HuH7 cells infected with influenza virus lacking NS1 compared to the WT virus (63), like the trend observed for *VILMIR*, suggesting that *VILMIR* could be regulated similarly as *BISPR* or other lncRNAs.

In A549 CRISPRi cells with *STAT1* knockdown, expression of *VILMIR* was not significantly affected. This was not expected since after suppression of host IFN responses (IAV vs. IAVdNS1 infection and pretreatment with Ruxolitinib before SARS-CoV-2 infection, Figure 2) there was a significant decrease in *STAT1* upregulation as well as *VILMIR* upregulation. In addition, the ChIP-seq analysis identified a STAT1 binding motif in the 5’ end region of *VILMIR* (Figure 1B), suggesting that STAT1 may regulate *VILMIR* expression. There are several possible explanations for these results. For example, as demonstrated by the RT-qPCR analysis of the *STAT1* KD cell lines, while basal expression of *STAT1* was reduced compared to the control cell line, *STAT1* transcription was not totally abolished in the *STAT1* KD lines. As activation and nuclear localization of STAT1 is regulated by its phosphorylation (62), it’s possible that a sufficient amount of activated STAT1 protein exists to induce certain ISGs such as *VILMIR*. Another explanation for this observation could be that *VILMIR* is not dependent upon STAT1. The decrease in *VILMIR* upregulation after IAV with an intact NS1 protein as well as after pretreatment with Ruxolitinib before SARS-CoV-2 infection (Figure 2) may be STAT1-independent, as NS1 is known to inhibit the IFN response through many different mechanisms (64), and Ruxolitinib specifically binds JAK1 and JAK2 (65). The JAK proteins can also phosphorylate other STAT complexes, such as STAT3 homodimers or STAT2:STAT3 heterodimers, which also have binding motifs within *VILMIR*. Additionally, while JAK/STAT represents the canonical IFN pathway, other non-canonical pathways exist such as MAP kinase (MAPK) and the phosphoinositide 3-kinases (PI3K)/mammalian target of rapamycin (mTOR) pathways that are activated by JAK1/TYK2 and have effects on ISG transcription (62). Further, the NF-κB and IRF pathways can induce ISGs independently of JAK signaling, such as RELB (subunit of NF-κB) and IRF1 which also had predicted binding motifs within *VILMIR*, and several lncRNAs have been found to be regulated by these pathways as well (66, 67). Future studies are needed to elucidate the exact mechanisms regulating *VILMIR* expression.

Airway epithelial cells are the first line of defense against respiratory viruses and play an important role in recruiting immune cells to fight infection (68), which is why our initial analysis of *VILMIR* started in epithelial cells. As we have demonstrated, however, *VILMIR* upregulation in response to SARS-CoV-2 infected clinical samples was observed in five immune cell types including macrophages, T cells, mDCs, natural killer cells, and B cells, as well as epithelial cells. In addition to this, it was upregulated after IFN-β treatment in T cell and monocyte cell lines. While many lncRNAs are cell type-specific, the upregulation of *VILMIR* in multiple immune cell types suggests it may play a broader role in immune responses. This also indicates that immune cells need to be investigated to determine the functional impact of *VILMIR in vivo*.

Through CRISPRi knockdown of *VILMIR* expression and RNA-seq based transcriptome profiling in A549 cells, we found that *VILMIR* KD broadly dampened the host transcriptional response to IFN-β treatment, i.e. genes that were upregulated or downregulated by IFN-β were still upregulated or downregulated, but with smaller fold changes. Interestingly, differentially expressed genes that were impacted by *VILMIR* knockdown were enriched in Interferon Signaling and JAK/STAT Signaling pathways, and several known ISGs such as *IFIT2* and *IFI44L* were also identified. While the overall magnitude of these changes we observed was relatively small, it may not necessarily reflect less significant biological effect. For example, while lncRNAs generally have low expression levels, several have been found to have significant outcomes on viral replication (69). In addition, the redundancy of ISGs presents a challenge in identifying antiviral functions specific to individual genes, as substantial antiviral response can still be induced in the absence of a specific ISG or even IFN signaling itself (70–72). Another possibility for small fold change differences at the transcription level is that *VILMIR* may function mainly by regulating at the post-transcriptional level, since *VILMIR* was found to be co-localized in both the nucleus and the cytoplasm. For example, lncRNA lnc-ISG20 can competitively bind microRNA 326, which in turn decreases the amount of microRNA 326 bound to ISG20 mRNA and therefore enhances the translation of ISG20 (15). LncRNA PYCARD-AS1, also distributed in both nucleus and cytoplasm, can interact with PYCARD mRNA in the cytoplasm and therefore inhibit ribosome assembly for PYCARD translation (73). Finally, while the knockdown of *VILMIR* expression was very efficient, it’s important to note that *VILMIR* transcription was not completely abolished, meaning that a low level of the transcript could still have functioned. Alternative approaches such as gene knockout or overexpression may be required for a full *VILMIR* functional perturbation.

In this study, we have identified a previously uncharacterized lncRNA, *VILMIR*, that shows a strong correlation to the host immune response to respiratory viral infection, and whose expression impacts the interferon response of host genes. We present this lncRNA as a novel ISG that should be further investigated in functional studies. Understanding the regulation of this gene in response to infection and how it functions within the host response may provide new insights into host-pathogen interactions.

## ACKNOWLEDGMENTS

The authors would like to thank Thomas Rowe from the Influenza Division of the Centers for Disease Control and Prevention for providing guidance on influenza propagation and titration. In addition, Barbara Sherry, Kenneth Adler, and members of the Peng lab from North Carolina State University College of Veterinary Medicine provided helpful feedback and critiques in manuscript preparation. We would also like to thank the UCSC CRISPR Core facility (RRID:SCR_021207) for their help and advice throughout this project. Finally, the authors would like to dedicate this manuscript to co-first author Ian Huntress, who passed away during the finalization of this manuscript, but whose computational work was foundational in the characterization of this lncRNA.

## DATA AVAILABILITY STATEMENT

The transcriptomic data discussed in this publication have been deposited in the Gene Expression Omnibus (GEO) https://www.ncbi.nlm.nih.gov/geo/ under accession number GSE261920.

